# Sarbecoviruses of British Horseshoe Bats; Sequence Variation and Epidemiology

**DOI:** 10.1101/2023.02.14.528476

**Authors:** Ternenge Apaa, Amy J. Withers, Ceri Staley, Adam Blanchard, Malcolm Bennett, Samantha Bremner-Harrison, Elizabeth A. Chadwick, Frank Hailer, Stephen W.R. Harrison, Mathew Loose, Fiona Mathews, Rachael Tarlinton

## Abstract

Horseshoe bats are the natural hosts of the *Sarbecovirus* subgenus that includes SARS-CoV-1 and 2. Despite the devastating impacts of the COVID-19 pandemic, there is still little known about the underlying epidemiology and virology of sarbecoviruses in their natural hosts, leaving large gaps in our pandemic preparedness. Here we describe the results of PCR testing for sarbecoviruses in the two horseshoe bat species (*Rhinolophus hipposideros* and *R. ferrumequinum*) present in Great Britain, collected in 2021-22 during the peak of COVID-19 pandemic. One hundred and ninety seven *R. hipposideros* samples from 33 roost sites and 277 *R. ferremequinum* samples from 20 roost sites were tested. No coronaviruses were detected in any samples from *R. ferrumequinum* whereas 44% and 56% of individual and pooled (respectively) faecal samples from *R. hipposideros* across multiple roost sites tested positive in a sarbecovirus-specific qPCR. Full genome sequences were generated from three of the positive samples (and partial genomes from two more) using Illumina RNAseq on unenriched samples. Phylogenetic analyses showed that the obtained sequences belong to the same monophyletic clade, with >95% similarity, as previously reported European isolates from *R. hipposideros*. The sequences differed in the presence or absence of accessory genes ORF 7b, 9b and 10. All lacked the furin cleavage site of SARS-CoV-2 spike gene and are therefore unlikely to be infective for humans. These results demonstrate a lack, or at least low incidence, of SARS-CoV-2 spill over from humans to susceptible GB bats, and confirm that sarbecovirus infection is widespread in *R. hipposideros*. Despite frequently sharing roost sites with *R. ferrumequinum*, no evidence of cross-species transmission was found.

## Background

The most widely accepted explanation for the origin of the SARS-CoV-2 pandemic is that it arose from animals held in the Wuhan market in late 2019 (Holmes, Goldstein et al. 2021). It has been proposed that masked palm civets (*Paguma larvatai*) acted a bridging host for SARS-CoV transmission from an unknown ancestral reservoir in a species of horseshoe bats to humans (Lau, Woo et al. 2005, Wang, Yan et al. 2005). Widespread onward transmission of SARS-CoV-2 from humans to other mammals has also occurred during the human pandemic, including large outbreaks in farmed mink (*Neovison vison*) (Eckstrand, Baldwin et al. 2021, Larsen, Fonager et al. 2021, Lu, Sikkema et al. 2021, Rabalski, Kosinski et al. 2021), and the establishment of a new reservoir in wild white-tailed deer (*Odocoileus virginianus*) in the USA (Hale, Dennis et al. 2022, Willgert, Didelot et al. 2022), and repeated infection (though without the establishment of endemic transmission) in domestic cats (*Felis catus*) (Hosie, Hofmann-Lehmann et al. 2021, Bienzle, Rousseau et al. 2022, Goletic, Goletic et al. 2022, Jairak, Chamsai et al. 2022). Repeated reports of transmission back to humans from these species have also occurred (Lu, Sikkema et al. 2021, Pickering, Lung et al. 2022, Sila, Sunghan et al. 2022) highlighting the risk of new animal reservoirs developing, potentially as sources of new viral variants that might evade vaccine-induced immunity. Sporadic cases have also been reported in domestic dogs (*Canis familiaris*), hamsters (*Mesocricetus auratu*s), cattle (*Bos Taurus)* and camels (*Camelus dromedarius)*, and a large range of primates, non-domestic felids, mustelids, hippos (*Hippopotamus amphibius*) and manatees (*Trichechus manatus*) (reviewed in (Pappas, Vokou et al. 2022).

The natural hosts of sarbecoviruses (SARS-like betacoronaviruses) are insectivorous horseshoe bats (represented by a single extant genus *Rhinolophus* within the family Rhinolophidae; superfamily Rhinolophoidea; sub-order Yinpterochiroptera) and the related old world roundleaf bats of the family Hipposideridae which share the same superfamily. There are around 180 known species of Rhinolophidae and Hipposideridae with ranges across Eurasia and Africa, and sarbecoviruses have been detected in at least 30 of them (Muylaert, Kingston et al. 2022). For species and sites where more extensive studies have been performed (generally SE Asia), it is apparent that species- and site-specific clustering of virus sequences occurs (Boni, Lemey et al. 2020, Lytras, Hughes et al. 2022). Geographical hotspots of both bat and viral species are apparent in SE Asia (Wong, Li et al. 2019, Muylaert, Kingston et al. 2022).

There are, in contrast, very few reports of sarbecoviruses in the sub-order Yangochiroptera: a small number of isolated reports in individual animals or pooled samples of Molossidae and Vespertilionidae families have been recorded in multispecies studies (Muylaert, Kingston et al. 2022, Seifert, Bai et al. 2022, Wang, Pan et al. 2022).

There is also evidence from multiple studies for considerable variation in both sarbecovirus carriage and sequences within host and viral lineages in bats, with implications for potential cross species transmission (Wang, Pan et al. 2022). While the exact determinants of what makes sarbecoviruses more likely to cross the species barrier into humans are not known, one clear limiting factor is the ability of the virus spike protein (the protein that binds to the host cell facilitating virus entry) to bind to the human ACE-2 protein (the receptor for SARS-CoV and SARS-CoV-2 in humans) (Conceicao, Thakur et al. 2020). Considerable mutation and antigenic variation is present in this spike protein in SARS-CoV-2 isolates and bat sarbecoviruses (Boni, Lemey et al. 2020, Li, Lai et al. 2021). In SARs-CoV-2 this protein also displays a distinctive cleavage site for the furin protease that is missing in many bat isolates and appears to be a critical factor in transmission and pathogenesis in humans (Chan and Zhan 2022). Complicating matters further, it is apparent that there is frequent viral recombination between different sarbecovirus lineages (Boni, Lemey et al. 2020, Lytras, Hughes et al. 2022).

Two species of horseshoe bats are present in Great Britain, the greater horseshoe bat (*Rhinolophus ferrumequinum)* and lesser horseshoe bat (*Rhinolophus hipposideros*), both with distributions largely restricted to southwest England and Wales. Neither are common at a national scale and both are regarded as rare in Europe being provided with considerable protection under the Habitats Directive and Eurobats Agreement (JNCC 2022, JNCC 2022). In Britain, populations comprise approximately 13,000 greater horseshoe bats and 50,000 lesser horseshoe bats, (Mathews, Kubasiewicz et al. 2018) but both are increasing over time, recovering from historically depressed levels. They are considered Least Concern on the British Red List, but the greater horseshoe bat is near threatened in Europe and there have been recent extinctions of populations in several countries (Temple and Terry 2007). Britain and Ireland form the extreme north-western end of their geographic ranges (only *R. hipposideros* is found in Ireland), which extends through central and southern Europe to Central Asia for lesser horseshoe bats and across Asia to Japan for greater horseshoe bats. The two species can share roost sites and both use a variety of roosts across the year, including maternity and hibernation sites (Russo 2022).

Sarbecoviruses have been reported previously from both bat species. Partial RDRP (Rnase dependent RNA polymerase) sequences have been described from Chinese, Italian and Bulgarian greater horseshoe bats (Drexler, Gloza-Rausch et al. 2010, Balboni, Palladini et al. 2011, Balboni, Gallina et al. 2012, Hu, Zeng et al. 2017, Muth, Corman et al. 2018) at a prevalence of 26-42% in European samples. Full sequences are also available for a number of Chinese, one Korean and one Russian greater horseshoe bat sarbecoviruses; the Asian sequences are phylogenetically very distinct from the European isolates (Lin, Wang et al. 2017, Alkhovsky, Lenshin et al. 2022, Sander, Moreira-Soto et al. 2022, Wu, Han et al. 2022).

Similarly, partial RDRP sequences have been reported in Slovenian and Polish lesser horseshoe bats (Rihtaric, Hostnik et al. 2010), at a prevalence of 31%, and Spanish lesser horseshoe bats at a prevalence of 7.1% (Muth, Corman et al. 2018). More recently full genome sequences from a Russian and British lesser horseshoe bat (Crook, Murphy et al. 2021, Alkhovsky, Lenshin et al. 2022) have also been reported. Resequencing of partial RDRP genes from the Spanish, Italian and Slovenian isolates (and the full genome sequences for the UK and Russian isolates) has demonstrated a lack of the furin cleavage site present in SARS-CoV-2 (Sander, Moreira-Soto et al. 2022). The European greater and lesser horseshoe bat sequences are also phylogenetically distinct from each other (Sander, Moreira-Soto et al. 2022) with the two Russian isolates displaying 59-95% amino acid similarity to each other depending on the gene (Alkhovsky, Lenshin et al. 2022).

In addition to their being hosts of various sarbecoviruses (Wong, Li et al. 2019), horseshoe bats have been identified as potentially susceptible to infection with SARS-CoV-2 (Cook, Grant et al. 2021, Common, Shadbolt et al. 2022, Logeot, Mauroy et al. 2022). This led us to screen British horseshoe bat populations, at the height of the SARS-CoV-2 pandemic in western Europe, for the presence of sarbecoviruses with the aim to confirm or rule out the establishment of SARS-CoV-2 circulation in these animals.

## Materials and methods

### Sample collection

A total of 517 bat samples (oronasal swabs, external swabs of the rectal region, and faecal samples), including 474 from horseshoe bats, (Table 1) were collected from roost sites or animals in care in the UK between April 2021 and February 2022 during which time human cases in the UK underwent successive alpha, delta and omicron SARS-CoV-2 waves (UKHSA 2022). Faecal samples included those collected from individuals in capture bags during routine population monitoring (during which sex and age were also recorded where possible) and pooled samples of fresh droppings collected from the floor of bat roosts. All samples were collected secondarily to regular licenced population monitoring efforts. Handling of animals was conducted under Natural England Licence 2022-61108-Sci-Sci (Mathews), and followed best practice guidelines for minimising the risk of human to bat SARS-CoV-2 transmission). Ethical approval was granted by the University of Nottingham School of Veterinary Medicine and Science Committee for Animal Research and Ethics (CARE), and the University of Sussex Animal Welfare and Ethical Review Board. Samples were preserved in RNAlater at room temperature and sent to the University of Nottingham where they were stored at -20 °C until RNA extraction. Samples were collected from five bat species from 54 sites: 197 from lesser horseshoe bats (*Rhinolophus hipposideros*) at 33 sites, 277 for greater horseshoe bats (*Rhinolophus ferrumequinum*) from 20 sites, 10 from common pipistrelles (*Pipistrellus pipistrellus*), 32 from Daubentons bats (*Myotis daubentonii*) and 1 from a Natterer’s bat (*Myotis nattereri*) (Table 1 and Supplementary information).

**Table 1:** Summary of bat species, sample type and positivity rates for Sarbecovirus E gene qPCR

| Bat Species | Type of sample | Number of samples | Sarbecovirus E gene qPCR |
| --- | --- | --- | --- |
| Lesser horseshoe | Oronasal swabs | 71 | 0 (0%) |
|  | Rectal swabs | 56 | 3 (5.4%) |
|  | Individual faeces samples | 39 | 17 (43.5%) |
|  | Pooled faeces samples | 31 | 17 (54.8%) |
| Greater horseshoe | Oronasal swabs | 90 | 0 |
|  | Rectal swabs | 69 | 0 |
|  | Individual faeces samples | 91 | 0 |
|  | Pooled faeces samples | 28 | 0 |
| Common pipistrelle | Oronasal swabs | 3 | 0 |
|  | Rectal swabs | - | - |
|  | Individual faeces samples | 7 | 0 |
|  | Pooled faeces samples | - | - |
| Natterer's | Oronasal swabs | 1 | 0 |
|  | Rectal swabs | - | - |
|  | Individual faeces samples | - | - |
|  | Pooled faeces samples | - | - |
| Daubenton's | Oronasal swabs | 5 | 0 |
|  | Rectal swabs | 8 | 0 |
|  | Individual faeces samples | 16 | 0 |
|  | Pooled faeces samples | - | - |

### RNA extraction, reverse transcriptase (RT) and RNA-dependent RNA polymerase (RDRP) gene coronaviruses generic conventional PCR and envelope gene sarbecovirus-specific real-time PCR

RNA extraction from bat faecal and oronasal swabs, and cell culture supernatant as positive control, was carried out using the Macherey-Nagel RNA tissue extraction kit as per manufacturer’s instructions. The Wuhan SARS-CoV-2 strain positive control sample used throughout this study was kindly donated by Dr Christopher Coleman (Division of Infection, Immunity and Microbes, School of Life Sciences, University of Nottingham, UK). RT was performed in two steps, using M-MLV-RT and random hexamer primers (Promega) as per manufacturer’s instructions. All cDNA products were stored at -20 oC for conventional PCR.

A generic pan-coronavirus PCR assay published by (Woo, Lau et al. 2005) was used to amplify a 440 bp fragment of the coronavirus RDRP gene using Q5® Hot Start High-Fidelity DNA Polymerase (New England Biolabs cat no: M0493S). PCR products were purified using the Nucleospin® extract II kit (Macherey-Nagel) according to manufacturer’s instructions and were Sanger sequenced (Eurofins UK).

Real-time PCR was carried out using the Promega GoTaq® Probe 1-Step RT-qPCR System (Promega) with Sarbecovirus-specific envelope gene primers from bat RNA samples as published by (Corman, Landt et al. 2020).

RNA and cDNA quality control was assessed via partial amplification of 148 bp of the bat mitochondrial cytochrome b gene using a published conventional PCR protocol (Lopez-Oceja, Gamarra et al. 2016).

### Next generation sequencing and genome analyses

RNA sequencing was performed by Novogene UK, using the Illumina NovaSeq 6000 platform. Quality filtering and trimming to remove adapters, duplicates and low quality reads was achieved using fastp v0.23.1 (Chen, Zhou et al. 2018). Kraken2 v2.1.2 was used for taxonomic classification reads against the Kraken2 viral Refseq database (Wood, Lu et al. 2019) (retrieved on 9th June 2022). Reads were assembled using the coronaSPAdes option in SPAdes genome assembler v3.15.4 (Meleshko, Hajirasouliha et al. 2021) using default parameters. While CheckV v1.0.1, a fully automated command-line pipeline, was used for identification and quality assessment of contigs, contigs were also queried against the NCBI custom BLASTN (v2.12.0) viral database (Altschul, Gish et al. 1990) (retrieved on 3rd July 2022) to confirm results from CheckV. Assembled contigs were indexed and contigs that were classified and assessed as complete bat Sarbecovirus genomes were extracted for downstream analysis using samtools v1.16.1 faidx option (Danecek, Bonfield et al. 2021). Individual reads for each sample were mapped back to identified contigs using Minimap2 (Li 2018), read coverage and depth were generated using samtools (Danecek, Bonfield et al. 2021). Assembled genomes were annotated in Geneious Prime® (v.2022.2.2) using NCBI coronavirus reference sequences and visualized for the presence of the structural, non-structural protein, accessory genes and to generate linear genome maps and data on individual gene location, composition, and nucleotide length. Reads were also mapped to the prototype European horseshoe bat (*Rhinolophus blasii*, Blasius horseshoe bat Bulgarian isolate) sarbecovirus reference genome (NC014470) using BBMap for variant calling, SNPs were viewed, and data exported in Geneious Prime® (v.2022.2.2).

### Phylogenetic analysis

Complete coronavirus genomes, extracted RDRP, spike, envelope and nucleocapsid nucleotide sequences from sarbecovirus genomes assembled in this study, and a total of 198 reference sarbecovirus genomes (including non-human sarbecovirus isolates, SARS-CoV and SARS-CoV-2) downloaded from NCBI, were aligned using mafft v7.490 (Katoh and Standley 2013). Maximum likelihood phylogenetic trees were reconstructed based on complete coronavirus genomes, and four different genes using IQ-TREE v2.0.7 (Minh, Schmidt et al. 2020), using 1000 bootstrap approximations following implementation of UFBoot2 within IQ-TREE v2.0.7 to evaluate branch support (Hoang, Chernomor et al. 2018). The ModelFinder option was included in command-line to select the best fitting nucleotide substitution model for phylogenetic reconstruction (Kalyaanamoorthy, Minh et al. 2017). Phylogenetic trees were visualized and annotated in FigTree v1.4.4 (https://github.com/rambaut/figtree/).

### Spike glycoprotein comparison, identification of furin cleavage site (FCS) and transmembrane protease serine 2 (TMPRSS2), receptor binding domain (RBD) homology modelling and structural analysis

To identify and compare the presence of FCS and TMPRSS2 between UK Coronavirus genomes and related CoVs, RBD spike proteins from the six UK Coronaviruses (five from this study and one reported by Crook et al 2020), SARS-CoV, SARS-CoV-2 and related Beta coronaviruses were aligned, viewed and pictures generated in Jalview (Procter, Carstairs et al. 2021). The model for the receptor binding domain (RBD) and hACE2 protein complex were constructed using SWISS-MODEL to assess the amino acid residues and structural differences between UK bat CoV, SARS-CoV, and SARS-CoV-2 using an RBD-hACE2 complex. The SWISS-MODEL template library was search for evolutionary related structures matching the target UK Coronavirus RBD (residues 321 11 515) and hACE2 (PDB: Q9BYF1) protein sequences using BLAST HHblits database. The most suitable template (PDB: 6vw1) with the highest selected Global Model Quality Estimate (GMQE) of 0.75 (range: 0 11 1) and the lowest average Quaternary Structure Quality Mean Estimate (QMEAN) of – 0.64 was selected to build models for the five UK Coronavirus RBD proteins generated from this study. The structural analysis and verification server (SAVES) was used to validate 3D structures, create Ramachandran plots, and classify amino acid residue torsion angles of each structure via implementation of PROCHECK AND ERRAT2 (Laskowski, MacArthur et al. 1993), respectively. In addition, receptor binding residues detection and binding energies of 3D model complex structures generated were calculated using PRODIGY (Vangone and Bonvin 2015, Honorato, Koukos et al. 2021). Visualization, superimposition, alignment, and generation of figures for presentation was done using UCSF Chimera (Pettersen, Goddard et al. 2004).

### Geospatial mapping and data analysis

Maps of roost sites positive and negative for coronavirus PCR were created in QGIS (version 3.28.2). to account for pooled and individual samples, map circle sizes were set to 5 proportional sizes as follows: <3 bats, 3-5 or one pooled sample, 5-10 (pools count as 3), 10-120 (pools count as 3) and >20 (pools count as 3). Chi-square test was used to evaluate potential differences in prevalence with sex and age-class (SPSS, Amos 28.0.0).

## Results

No samples tested positive in the pan-coronavirus assay, and no oronasal swabs tested positive in the sarbecovirus-specific PCR assay for either species, However, numerous lesser horseshoe bat faecal samples tested positive with the Sarbecovirus qPCR assay (Table 1). Within the five roosts from which >5 individual faecal samples and/or rectal swabs were collected, all yielded at least one positive sample, and the percentage positivity ranged from 5% to 25% (mean at individual level across those sites, 16/100 individuals, 16%, 95% CI (Wilson’s) 10.0%-24.6%). Twenty-two sites yielded only single, pooled samples, of which 10 (45%; 95% CI (Wilson’s) 27.0%-65.3%) were positive (Supplementary information). The locations and infection status of lesser horseshoe bat roost sites yielding either >3 individual faecal samples or at least 1 pooled faecal sample are shown in Figure 1; of these, 21/31 (68%; 95% CI (Wilson’s) 50.0%-81.4%) roosts yielded at least one positive sample.

**Figure 1:**
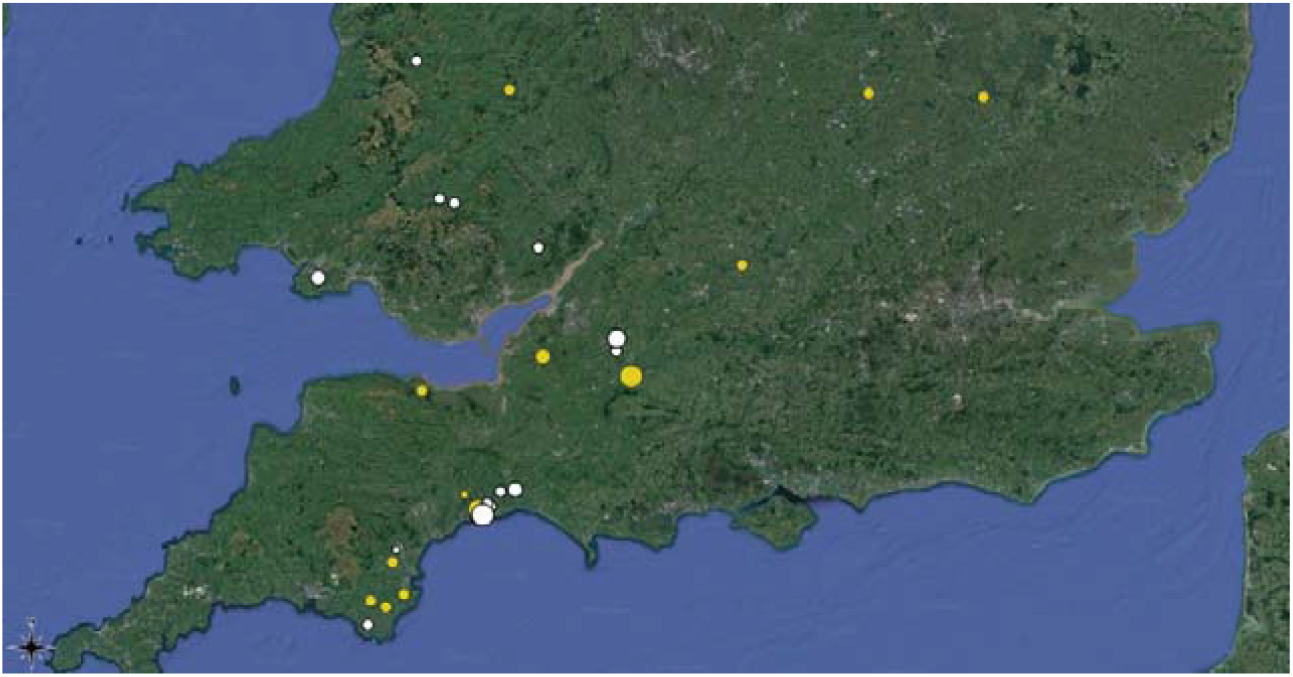
Roost sites of *Sarbecovirus* detected by PCR in faeces from lesser horseshoe bats (*Rhinolophus hipposideros)*, Yellow=negative, White= positive. Circle sizes are proportional to number of samples from the site.

Age and sex were recorded for 41 and 37 animals, respectively, when individual faecal samples were collected and examined; among this subset of animals tested, there were no significant (chi-square) differences in the frequency of positive samples by age (19.5% adult, 8% juvenile) or sex (12.5% female, 21.9% male).

### Taxonomic classification, genome assembly

Taxonomic classification using Kraken2 identified reads assigned to other viral operational taxonomic units, however, only reads classified to the Coronaviridae viral family are reported in this study. Paired end reads were generated from RNA extracted from six lesser horseshoe bat faecal samples. Between 98 to 99% of reads were unclassified. The percentage of classified viral reads from the datasets examined varied from 0.49 11 1.18%, while the total percentage of classified viral reads assigned to the family Coronaviridae varied from 0.01 11 63.58%. Three samples including RhGB04, RhGB05 and RhGB06 recorded approximately 50% of classified viral reads assigned to Coronaviridae (Supplementary information).

De novo assembly of datasets from lesser horseshoe bats yielded five genomes made up of single contigs. The length of the assembled genomes varied from 28.2 11 30.6 kb with one short contig of 12.6k. CheckV analysis demonstrated that all the assembled contigs were closely related to beta coronaviruses, specifically the subgenus Sarbecovirus. Sarbecovirus genomes identified were assessed to have 94 11 100% (4/5) and 42% (1/%) quality/completeness, all genomes had 0% contamination except for RhGB05 with 3.17% contamination. Overall, all the assembled genomes shared 94.5-97.7% average amino acid identity with the reference genome (RhGB01 MW1957) identified by CheckV (Supplementary information).

Assessment of reads mapped to the assembled genomes demonstrated over 7000, 4000, 302,000, 420,000 and 500,000 reads mapped to RhGB02, RhGB03, RhGB04, RhGB05 and RhGB06, respectively. The genome assembled from RhGB03 had the least average mean depth of 22X and was missing ORF10 and the 3’ UTR, while RhGB04 (partial genome of 12.6 kb missing the 5’ end of ORF 1ab) recorded the highest mean depth coverage of 3446X (Supplementary information).

### Genome annotation and organization of British bat CoVs

Genome annotation in Geneious Prime using NCBI SARS-CoV-2 and SARS-related reference sequences, confirmed a similar genome organization to the UK lesser horseshoe bat reference genome available from the NCBI database (Figure 2). Sarbecovirus genomes consist of a leader sequence (5’UTR), followed by ORF1ab gene with sixteen non-structural proteins (nsp1 11 16) making up about 2/3 of the viral genome. The assembled sarbecoviruses were made up of four major structural proteins including the spike (S), membrane (M), envelope (E) and nucleocapsid (N) proteins. While accessory proteins including ORF3b, 6a, 6b, 7a and variable ORFs recognisable for 7b, 9b and 10 were reported within the 3’ region interspaced between the major structural proteins (Figure 2).

**Figure 2:**
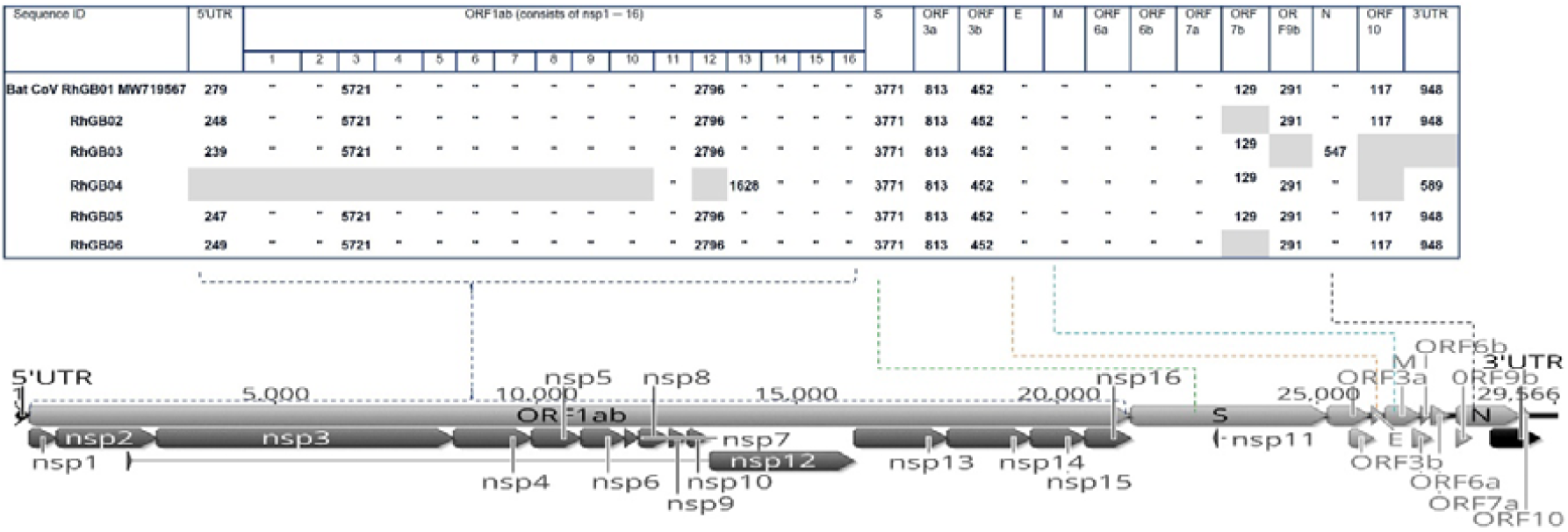
Genome organisation of lesser horseshoe bat Sarbecovirus sequences derived from this study and the reference UK genome RhGB001. Missing genes are shown in grey, lengths of nucleic acid segments are listed for each gene, LTR and isolate.

### Variant calling

Variant calling following mapping of raw reads to the bat Bulgarian Sarbecovirus reference genome (Bat CoV BM48-31) available in Europe, demonstrated the presence of a series of single nucleotide polymorphism (SNPs) characterized by amino acid substitution distributed throughout the entire length of the genome. However, only a total of 22 unique SNPs were identified within the spike glycoprotein (Supplementary information) with minimum read coverage ≥ 10X, mapped quality ≥ 10% and variant frequency > 95% from the four full genomes assembled (Supplementary information for details). Overall, 15/22 SNPs resulting in amino acid substitution were found in RhGB02, 12/22 SNPs identified in RhGB03(12/22) and 8/22 SNPs were reported from both RhGB05 and RhGB06 with the highest read-depth coverage. One unique SNP at coding sequence position 3641 (V to A) was found in all four assembled genomes, and 5/22 unique SNPs, including changes at position 2399 (SR to KR), 2766 (D to E), 2781 (TT to TA), 3208 (E to K), and 3268 (I to V), in at least 3/4 of the assembled genomes. The position of these SNPs coincided with location of Sarbecovirus SD-1 and SD-2 subdomains, S1/S2 cleavage region and the S2 fusion subunit (Supplementary information).

### Phylogenetic analysis

Results from the maximum likelihood phylogenetic trees drawn using complete coronavirus genomes (Supplementary information), spike glycoprotein (Figure 3) RDRP envelope and nucleocapsid (Supplementary information) nucleotide sequences showed that the Sarbecovirus genomes assembled from UK lesser horseshoe bats belong to the same monophyletic clade as the published European horseshoe bat Sarbecovirus genomes (RhGB01, Khosta1 and 2 and BM48-31), and are more similar to the small number of African horseshoe bat sarbecoviruses than to any Asian isolates, even those from the same host species.

**Figure 3:**
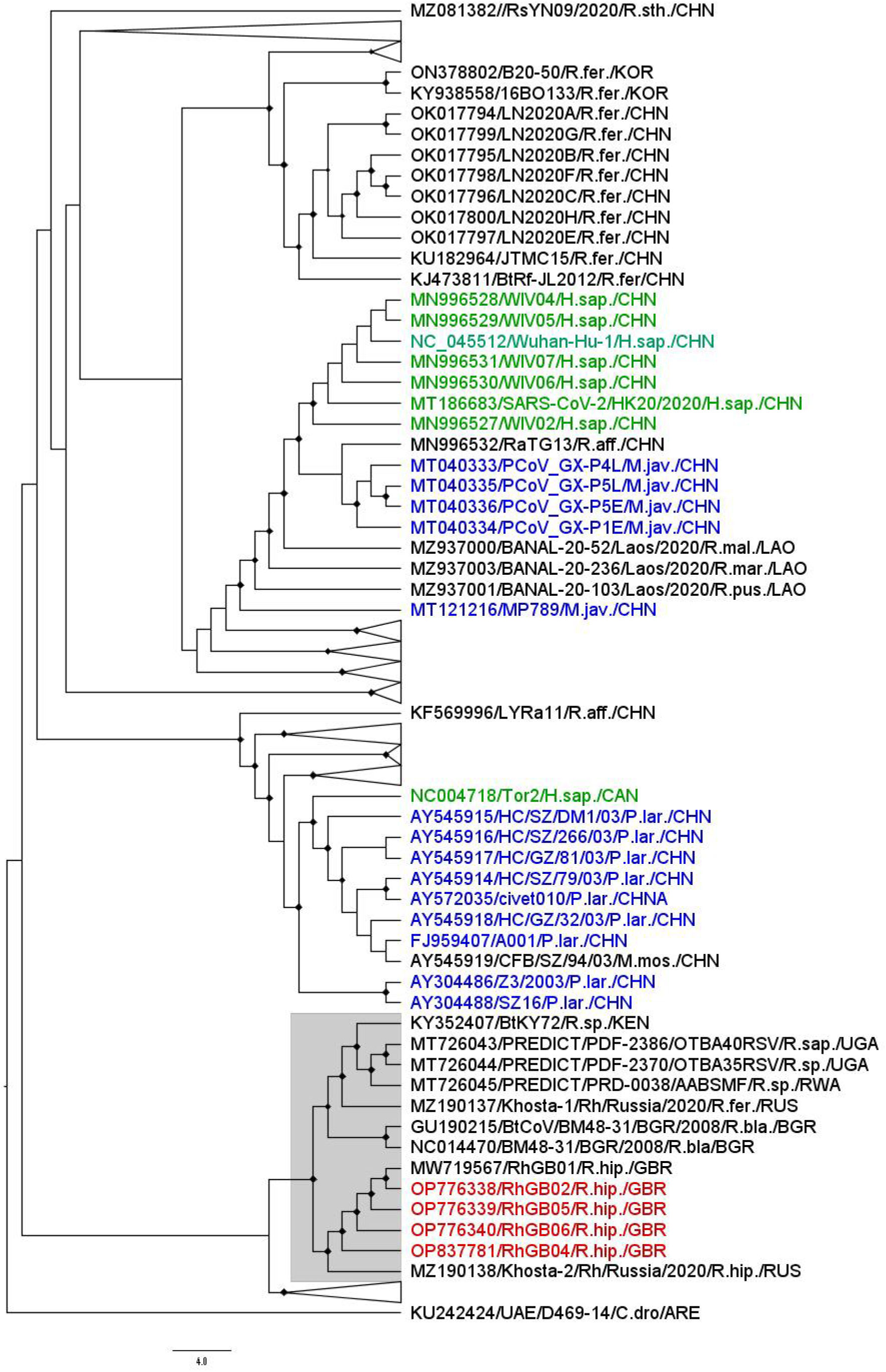
Maximum likelihood phylogenetic tree of S gene nucleic acid constructed with 1000 bootstrap approximation, rooted on the MERs coronavirus reference sequence. One hundred and ninety eight non-human Sarbecovirus genomes and reference sequences for major variants of SARS-CoV and SARS-CoV-2 were included (non-human SARS-CoV-2 isolates were not included). Clades of Asian bat coronavirus sequences (apart from greater horseshoe bat sequences) have been collapsed for clarity (represented as broad triangles). Red=isolates from this study, Green=human isolates, blue=isolates from other mammals. Sequences are named with Genbank ID, name from original study species of origin (eg R.hip= *Rhinolophus hipposideros*) and country of origin (eg GBR= Great Britain)

### Spike glycoprotein comparison, identification of furin cleavage site (FCS) and Transmembrane protease serine 2 (TMPRSS2), receptor binding domain (RBD) homology modelling and structural analysis

All the UK bat coronavruses reported in this study shared the same spike glycoprotein amino acid length of 1256 aa with the only available complete bat SARS-like coronavirus genome reported from UK lesser horseshoe bats (Crook et al., 2021). The S1 RBD of UK bat coronaviruses reported from this study and the previously reported RhGB01 (Crook, Murphy et al. 2021) consisted of 220 amino acids, the RBM (receptor binding motif) of UK lesser horseshoe bat coronaviruses reported from this study have the same amino acid length (71 aa) as RhGB01.

The RBD (receptor binding domain) pairwise alignment percentage identity estimation (figure 4B) between the reported UK bat coronaviruses and selected Beta coronavirus reference sequences revealed an estimated percentage identity of 68% with SARS-CoV and 65 11 67% with SARS-CoV-2 related viruses. The RBD of all five UK bat coronaviruses reported from this study shared ≥ 95% amino acid homology with the RhGB01 reported by Crook, Murphy et al. (2021), and 76% amino acid homology with the Bulgarian horseshoe bat sarbecovirus (BM48-31) reported by Drexler, Gloza-Rausch et al. (2010). Further assessment of the S1 and S2 regions of UK bat coronaviruses RBDs showed the absence of a furin or S1/S2 cleavage site (figure 4A), while demonstrating the presence of host transmembrane serine protease 2 (TMPRSS2) or S2 cleavage site from reported UK bat CoVs (figure 4A).

**Figure 4.**
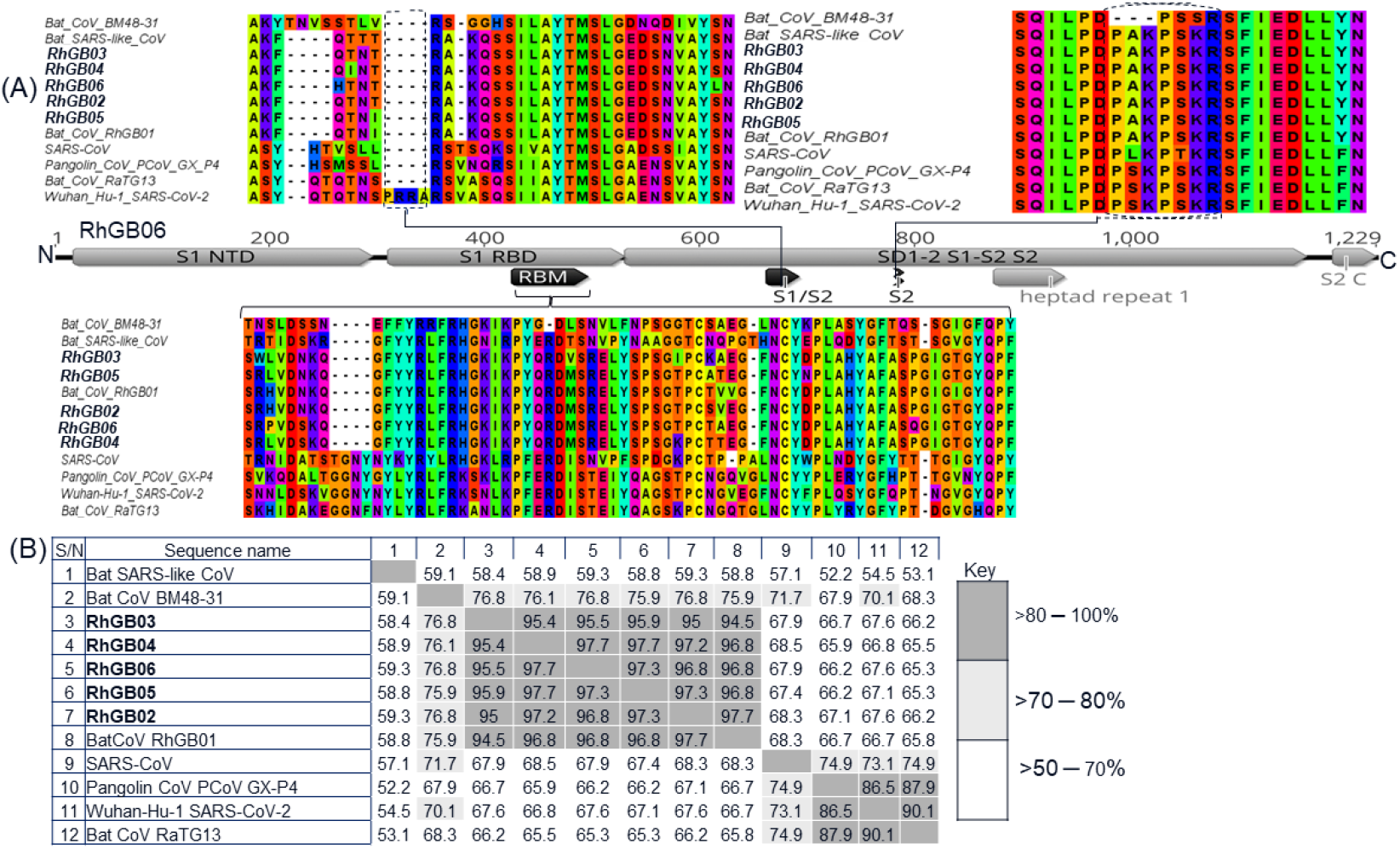
British bat Sarbecovirus (in bold text) spike glycoprotein organization and sites of interest. (A) The presence of conserved S2 or TMPRSS2 cleavage site (top right) and the absence of furin cleavage site (S1/S2 top left). The FCS consisted of four amino acids (PRRA) identified to be present in only SARS-CoV-2. Below these two is the British bat CoVs spike glycoprotein linear map. (B) Estimated percentage pairwise alignment identity heatmap following alignment of RBD proteins from sarbecovirus reference genomes and those reported in this study.

Comparative homology modelling of this study’s sarbecoviruses to examine the interaction between RBD and the human ACE2, yielded five UK bat CoV 3D models sharing ≥ 92% identity and 3.0Å root-mean-square deviation (RMSD) of C-atoms with the SARS-CoV-2 (PDB: 6vw1) X-ray crystal template structure (figure 5A and B). The overall quality factor of 3D models was calculated through validation with ERRAT2 and Ramachandran plots (Supplementary information) as ≥ 96 for all RBD and hACE2 3D structures constructed. Amino acid residues rejected at 95% and 99% confidence intervals and those known to be characteristic for the interaction between SARS-CoV-2 and hACE2 are highlighted along the RBD proteins in Figure 5. Ramachandran plots generated following ERRAT2 validation demonstrated that 91% of residues were in the most favoured regions, 8.7% in additional allowed regions and 0.3% were found to be in the generously allowed regions (Supplementary information).

**Figure 5.**
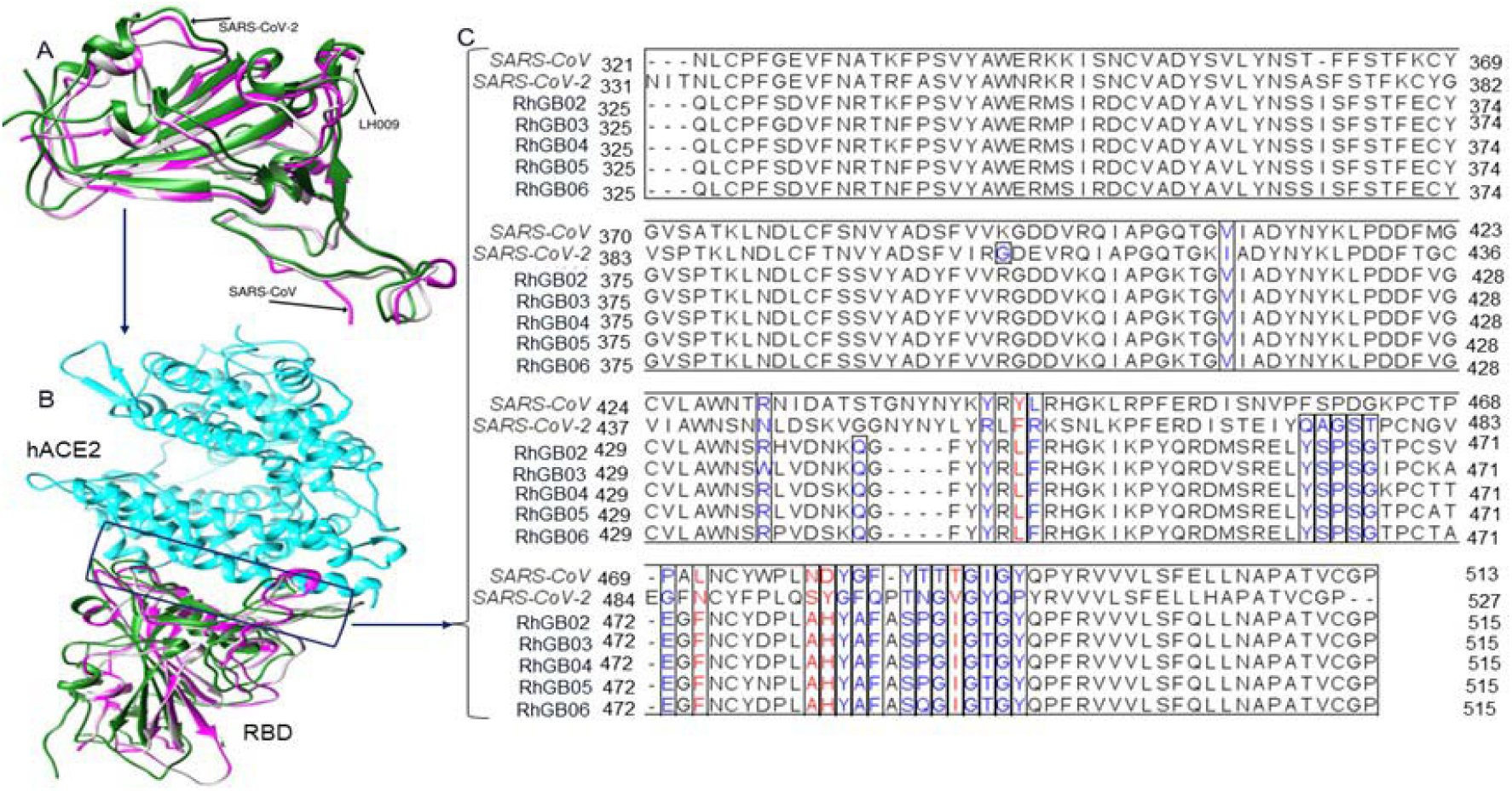
Protein-protein homology complex model of British bat and related sarbecoviruses (A) Superimposed RBD 3D complexes from UK bat CoVs (gray LH009), SARS-CoV (magenta), and SARS-CoV-2 (green). (B) Superimposed model between RBDs of UK bat CoVs (gray), SARS-CoV (magenta) and SARS-CoV-2 (green) and hACE2 (cyan). (C) Sequence alignment and comparison of S1 RBD and RBM of UK bat CoVs, SARS-CoV, and SARS-CoV-2. RBM for UK bat CoVs is positioned between 426 aa to 496 aa. Amino acid residues predicted to interact with hACE2 are shown in blue text, amino acid residues in red text are the critical residues within the RBM previously reported to play key role in cross-species transmission.

Prediction of amino acid residues using Prodigy showed that amino acid residues at interface with hACE2 receptor in British bat Sarbecovirus spike glycoproteins can be located at positions V409, R431, Q437, Y441, L443, F444, Y461, S462, P463, S464, G465, E471, F473, Y482, A483, F484, S486, P487, G488, I489, G490, T491, and Y493; as reported previously (Crook, Murphy et al. 2021). RBM of these sarbecoviruses is located between amino acid position 426 aa to 496 aa of the spike protein, coinciding with the location of hACE2 amino acid residues. In addition, RBD comparative analysis revealed the presence of critical amino acid residues in British bat sarbecoviruses (L443, L443, F473, A480, H481 and I489) previously reported to play a key role in cross-species transmission of SARS-like CoVs (Figure 5).

## Discussion

This study found no detectable SARS-CoV-2 in British bats, in particular none in horseshoe bats which might be expected to be most at risk of infection. This is similar to the findings of other studies of European bats during the COVID-19 pandemic, none of which demonstrated SARS-CoV-2 infection including from, three common pipistrelles in the UK (Jones, Bell et al. 2022), 503 samples from 20 bat species including 58 lesser horseshoe bats in Poland (Orłowska, Smreczak et al. 2022), 197 samples from five bat species including 82 samples from lesser horseshoe bats, 104 from greater horseshoe bats and five from Mediterranean horseshoe bats (*Rhinolophus euyale)* from Sochi in Russia (Alkhovsky, Lenshin et al. 2022) and 76 lesser horseshoe bats in the UK (Crook, Murphy et al. 2021).

The pan-coronavirus screening assay used in this study (Woo, Lau et al. 2005) is known to be insensitive and may miss some coronavirus strains due to sequence mismatch for example, it is not a good match for the known UK isolate MW719567, RhGB01. The Sarbecovirus E gene qPCR (Corman, Landt et al. 2020) was designed to detect all known sarbecoviruses and is in common use in human SARS-CoV-2 diagnostics due to the relative stability of the E gene in SARS isolates and sarbecoviruses in general. Serial dilution of the SARS-CoV-2 positive control in this study (data not shown) found that the E gene qPCR was approximately a 102 times more sensitive than the pan coronavirus screening assay, though SARS-CoV-2 is routinely detected by both assays. This difference in sensitivity of the two assays may explain the discrepancies in results between the two screens (CT values for the E gene qPCR ranged between 22 and 40). It is therefore possible that other coronaviruses, in particular non-sarbecoviruses are present in these sample but were not detected.

British lesser horseshoe bats were, however, found to be frequently infected with a Sarbecovirus similar to that described previously in this species (Rihtaric, Hostnik et al. 2010, Crook, Murphy et al. 2021, Alkhovsky, Lenshin et al. 2022, Orłowska, Smreczak et al. 2022, Sander, Moreira-Soto et al. 2022) and distinct from sarbecoviruses previously described in greater horseshoe bats in Bulgaria (Muth, Corman et al. 2018) and Russia (Alkhovsky, Lenshin et al. 2022). Although the sampling strategy in this study, based on opportunistic sampling linked to bat survey and conservation studies, did not allow a prevalence to be calculated, the frequency at which the virus was detected within and between populations suggests a prevalence not dissimilar to that found in Poland of around 30% (Orłowska, Smreczak et al. 2022). That study, unlike this, detected virus in oronasal swabs as well as faeces, and this may reflect the smaller amount of material collected on swabs in this study (rectal swabs were also less frequently positive in the qPCR assay than faecal samples). The sites sampled reflected the distribution of horseshoe bats in Great Britain, and infection was clearly common within and between populations of lesser horseshoe bats in this study, with no clear sex or age differences in likelihood of shedding. This high frequency of shedding and widespread distribution geographically and demographically, likely indicates either persistent infection and excretion, or frequent reinfection, of the gastrointestinal tract as has been reported for many other coronaviruses, such as those of cats, chickens and pigs (Quinteros, Noormohammadi et al. 2022, Thayer, Gogolski et al. 2022, Zhang, Chen et al. 2022).

Many of the roosts of lesser horseshoes bats in this study were shared with greater horseshoe bats, yet none of the latter were shedding detectable virus. This suggests lack of cross-species transmission and that the virus is relatively host-species restricted. This contrasts with several studies of SE Asian horseshoe bat species and their sarbecoviruses (Latinne, Hu et al. 2020, Cappelle, Furey et al. 2021, Zhou, Ji et al. 2021), which found strong evidence of cross-species transmission of coronaviruses. Lesser horseshoe bats were primarily sampled in the summer due to the greater sensitivity of this species to disturbance, with greater horseshoe bats sampled across both summer and winter (53/277). The numbers of greater horseshoe bats in this study should have been adequate to detect, if it were present, the virus previously detected in Italian, Bulgarian and Russian greater horseshoe bats at the prevalence rates of 26-42% recorded in those studies (Drexler, Gloza-Rausch et al. 2010, Balboni, Palladini et al. 2011, Balboni, Gallina et al. 2012, Muth, Corman et al. 2018, Alkhovsky, Lenshin et al. 2022) and the assay should be able to detect the Russian isolate (the only one for which an E gene is available). It is not clear why no sarbecoviruses were detected in these animals, though annual variation in viral loads is possible as is a “founder effect” in an island population on the edge of the host’s geographic range.

The apparent host specificity of the European lesser horseshoe bat Sarbecovirus, combined with its lack of the furin cleavage site thought to be critical for human spread of SARS-Cov-2, indicate that these viruses are likely of low potential for zoonotic transmission, although modelling studies (Sander, Moreira-Soto et al. 2022) have indicated that these viruses could acquire such features with minimal mutation and that these viruses could potentially bind to the human ACE-2 receptor (Crook, Murphy et al. 2021).

The sarbecoviruses found in European lesser horseshoe bats cluster monophyletically and probably represent a distinct genus of Sarbecovirus. Recombination analysis with RDP5 (data not shown) indicates that these viruses are not recombinants.

Some variation in the presence of the accessory genes 7b, 9b and 10 was found amongst the isolates studied. These genes are not essential for viral replication, but are modulators of the host’s innate immune system response and as such can affect strain pathogenicity (Fang, Fang et al. 2021, Forni, Cagliani et al. 2022, McGrath, Xue et al. 2022). Plasticity in accessory gene complement has also been evident in different SARs-CoV-2 isolates and appears to be a feature of Sarbecovirus isolate variability (Forni, Cagliani et al. 2022, McGrath, Xue et al. 2022).

We do not have any indication if infection with these viruses have any adverse affects on their hosts or whether excretion patterns vary with age or reproductive status. This type of work is mature in relation to other bat-borne viruses, for example Hendra virus (a member of the Paramyxoviridae family) in bats of the genus Pteropus (flying foxes), for which it is clear that virus excretion is associated with maternity roosts with large numbers of birthing and juvenile animals and that virus excretion peaks in times of nutritional stress (Eby, Peel et al. 2022). Such work is however in its infancy with horseshoe bat sarbecoviruses, although some longitudinal studies in SE Asia hint at a summer/maternity roost excretion pattern (Cappelle, Furey et al. 2021).

A better understanding of the ecology of horseshoe bat viruses requires further and longer term studies of sarbecoviruses in their natural hosts. The samples with the highest detection rates in this study were pooled faecal samples from bat roosts so this sample type may provide the most reliable method of detection of virus in a roost site as well as being the most convenient, causing least disruption to bat colonies and presenting least threat of cross-species transmission between bats or between bats and humans.

Overall, this study provides several critical pieces of data in the overall picture of sarbecoviruses in their natural hosts. The growing picture is one of relative stability in terms of viral diversity and cross-species transmission potential in European horseshoe bat species, in contrast to higher diversity and cross species sharing of isolates in SE Asia, potentially related to the greater diversity and number of horseshoe bat species in the latter geographic region. This likely partly explains the repeated cross-species spill over of sarbecoviruses with an ancestral origin in bats, into civets, pangolins and humans in SE Asia, though human behaviour and wildlife/human interactions, habitat disruption and farming of wildlife for meat are also likely to be contributing factors (Holmes, Goldstein et al. 2021). Ongoing work with these European sarbecoviruses remains to determine the epidemiology, temporal changes in viral prevalence and sequence and effects of the virus in its natural host with a real need for longitudinal studies of roost sites across the host range and spatial comparisons of sequences and prevalence.

## Ethical approval

Ethical approval was granted by the University of Nottingham School of Veterinary Medicine and Science Committee for Animal Research and Ethics (CARE) and the Animal Welfare and Ethical Review Board of the University of Sussex.

## Acknowledgements

We are grateful for the help of many licensed bat workers who assisted in the collection of samples for this project, particularly Devon Bat Research and Conservation Group, Wiltshire Bat Group, and the Vincent Wildlife Trust. We also appreciate constructive discussions with the Bat Conservation Trust that informed this project.

