## Supplementary Information for "Sarbecoviruses of British Horseshoe Bats; Sequence Variation and Epidemiology"

Supplementary Table 1 Roost site locations, species and % positivity for E gene qPCR (two samples did not have an identifiable GPS location from the submitter. Sites have been anonymised to 1 decimal place (11.1km)

| Roost site location (GPS coordinates) | Sarbecovirus E gene qPCR results | *Rhinolophus hipposideros*  (% positive for E gene qPCR) | *Rhinolophus ferrumequinum* | *Pipistrellus* *pipistrellus* | *Myotis daubentonii* | *Myotis nattereri* |
| --- | --- | --- | --- | --- | --- | --- |
| Site 1 | negative | 1 | 7 |  |  |  |
|  | positive | 1 (50%) |  |  |  |  |
|  | total | 2 | 7 |  |  |  |
| 50.5, -3.6 | negative | 5 | 3 |  | 2 | 1 |
|  | positive | 1 (16%) |  |  |  |  |
|  |  | 6 | 3 |  | 2 | 1 |
| 51.3, -2.2 | negative | 30 | 16 |  | 20 |  |
|  | positive | 1 (3%) |  |  |  |  |
|  | total | 31 | 16 |  | 20 |  |
| 50.7, - 3.0 | negative |  | 1 |  |  |  |
|  | positive | 1 (100%) |  |  |  |  |
|  | total | 1 | 1 |  |  |  |
| 50.6, -3.6 | negative | 21 | 69 |  | 8 |  |
|  | positive | 4 (16%) |  |  |  |  |
|  | total | 25 | 69 |  | 8 |  |
| 50.6, -3.1 | negative | 64 | 56 |  |  |  |
|  | positive | 9 (14%) |  |  |  |  |
|  | total | 73 | 56 |  |  |  |
| 50.6, -3.1 | negative | 6 |  |  |  |  |
|  | positive | 2 (25%) |  |  |  |  |
|  | total | 8 |  |  |  |  |
| 51.6, -4.1 | negative | 1 |  |  |  |  |
|  | positive | 2 (66%) |  |  |  |  |
|  | total | 3 |  |  |  |  |
| 50.7, -2.9 | negative | 7 | 2 |  |  |  |
|  | positive | 2 (22%) |  |  |  |  |
|  | total | 9 | 2 |  |  |  |
| 52.3, -0.6 | negative | 1 |  |  |  |  |
|  | positive |  |  |  |  |  |
|  | total | 1 |  |  |  |  |
| [52.6, -2.8](http://www.nearby.org.uk/coord.cgi?p=SJ+40290+01584#llSJ4029001584) | negative |  |  |  |  |  |
|  | positive | 1 (100%) |  |  |  |  |
|  | total | 1 |  |  |  |  |
| 51.3, -2.2 | negative | 4 |  |  |  |  |
|  | positive | 1 |  |  |  |  |
|  | total | 5 (20%) |  |  |  |  |
| 50.7, -3.1 | negative | 3 |  |  |  |  |
|  | positive | 1 (25%) |  |  |  |  |
|  | total | 4 |  |  |  |  |
| 50.6, -3.1 | negative | 3 |  |  |  |  |
|  | positive |  |  |  |  |  |
|  | total | 3 |  |  |  |  |
| 50.7, -3.2 | negative | 4 |  |  |  |  |
|  | positive |  |  |  |  |  |
|  | total | 4 |  |  |  |  |
| 51.3, -2.7 | negative | 2 |  |  |  |  |
|  | positive | 1 (33%) |  |  |  |  |
|  |  | 3 |  |  |  |  |
| 51.9, -3.3 | negative |  |  |  |  |  |
|  | positive | 1 |  |  |  |  |
|  | total | 1 |  |  |  |  |
| 52.4, -3.5 | negative |  |  |  |  |  |
|  | positive | 1 (100%) |  |  |  |  |
|  | Total | 1 |  |  |  |  |
| 51.7, -2.7 | positive | 1 |  |  |  |  |
|  | total | 1 |  |  |  |  |
| [52.3 -2.9](http://www.nearby.org.uk/coord.cgi?p=SO+33031+74358&f=full#llSO3303174358) | negative | 1 |  |  |  |  |
|  | Positive |  |  |  |  |  |
|  | total | 1 |  |  |  |  |
| 51.6, -1.5 | negative  positive | 1 |  |  |  |  |
|  | total | 1 |  |  |  |  |
| 51.2, -1.0 | negative | 1 |  |  |  |  |
|  | positive |  |  |  |  |  |
|  | total | 1 |  |  |  |  |
| 51.6, -2.1 | negative | 1 |  |  |  |  |
|  | positive |  |  |  |  |  |
|  |  | 1 |  |  |  |  |
| 50.5, -3.6 | negative | 1 |  |  |  |  |
|  | positive |  |  |  |  |  |
|  | total | 1 |  |  |  |  |
| 50.3, -3.6 | negative | 1 |  |  |  |  |
|  | positive |  |  |  |  |  |
|  | total | 1 |  |  |  |  |
| 50.3, -3.7 | negative | 1 |  |  |  |  |
|  | positive |  |  |  |  |  |
|  | total | 1 |  |  |  |  |
| 50.2, -3.8 | negative |  |  |  |  |  |
|  | positive | 1 |  |  |  |  |
|  | total | 1 |  |  |  |  |
| 50.3, -3.5 | negative | 1 |  |  |  |  |
|  | positive |  |  |  |  |  |
|  | total | 1 |  |  |  |  |
| 50.6, -3.1 | negative | 1 |  |  |  |  |
|  | positive |  |  |  |  |  |
|  | total | 1 |  |  |  |  |
| 50.7, -3.0 | negative |  |  |  |  |  |
|  | positive | 1 (100%) |  |  |  |  |
|  | total | 1 |  |  |  |  |
| 50.7, -3.0 | negative |  | 1 |  |  |  |
|  | positive | 1 (100%) |  |  |  |  |
|  | total | 1 | 1 |  |  |  |
| 50.7, -3.0 | negative |  | 1 |  |  |  |
|  | positive | 1 (100%) |  |  |  |  |
|  | total | 1 |  |  |  |  |
| [51.2, -2.2](http://www.nearby.org.uk/coord.cgi?p=ST859498&f=full#llST8590049800) | negative | 1 |  |  |  |  |
|  | positive |  |  |  |  |  |
|  | total | 1 |  |  |  |  |
| 51.3, -2.7 | negative | 1 |  |  |  |  |
|  | positive |  |  |  |  |  |
|  | total | 1 |  |  |  |  |
| 50.4, -3.7 | negative |  | 21 |  |  |  |
|  | positive |  |  |  |  |  |
|  | total |  | 21 |  |  |  |
| 50.8, -2.1 | negative |  | 26 |  |  |  |
|  | positive |  |  |  |  |  |
|  | total |  | 26 |  |  |  |
| 51.3, -2.2 | negative |  | 41 |  |  |  |
|  | positive |  |  |  |  |  |
|  | total |  | 41 |  |  |  |
| 50.6, -3.1 | negative |  | 2 |  |  |  |
|  | positive |  |  |  |  |  |
|  | total |  | 2 |  |  |  |
| 51.3, -2.2 | negative |  | 1 |  |  |  |
|  | positive |  |  |  |  |  |
|  | total |  | 1 |  |  |  |
| [50.3 -3.7](http://www.nearby.org.uk/coord.cgi?p=SX+736+526&f=full#llSX7360052600) | negative |  | 1 |  |  |  |
|  | positive |  |  |  |  |  |
|  | total |  | 1 |  |  |  |
| Site 2 | negative |  | 2 |  |  |  |
|  | positive |  |  |  |  |  |
|  | total |  | 2 |  |  |  |
| 50.3, -3.9 | negative |  | 1 |  |  |  |
|  | positive |  |  |  |  |  |
|  | total |  | 1 |  |  |  |
| 50.3, -3.5 | negative |  | 1 |  |  |  |
|  | positive |  |  |  |  |  |
|  | total |  | 1 |  |  |  |
| 50.3,-3.7 | negative |  | 1 |  |  |  |
|  | positive |  |  |  |  |  |
|  | total |  | 1 |  |  |  |
| 50.3, -3.4 | negative |  | 1 |  |  |  |
|  | positive |  |  |  |  |  |
|  | total |  | 1 |  |  |  |
| 50.6 -3.1 | Negative |  | 1 |  |  |  |
|  | positive |  |  |  |  |  |
|  | total |  | 1 |  |  |  |
| 50.7 -3.0 | negative |  | 3 | 5 |  |  |
|  | positive |  |  |  |  |  |
|  | total |  | 3 | 5 |  |  |
| 52.0, -3.7 | negative |  | 1 |  |  |  |
|  | positive |  |  |  |  |  |
|  | total |  | 1 |  |  |  |
| 50.7, -3.1 | negative |  | 1 |  |  |  |
|  | positive |  |  |  |  |  |
|  | total |  | 1 |  |  |  |
| 50.3, -4.0 | negative |  | 3 |  |  |  |
|  | positive |  |  |  |  |  |
|  | total |  | 3 |  |  |  |
| [51.2, -2.2](http://www.nearby.org.uk/coord.cgi?p=ST859498&f=full#llST8590049800) | negative |  | 2 |  |  |  |
|  | positive |  |  |  |  |  |
|  | total |  | 2 |  |  |  |
| 50.3, -4.0 | negative |  | 12 |  |  |  |
|  | positive |  |  |  |  |  |
|  | total |  | 12 |  |  |  |
| 49.9, -5.1 | negative |  |  | 4 |  |  |
|  | positive |  |  |  |  |  |
|  | total |  |  | 4 |  |  |
| [51.1,1.2](http://www.nearby.org.uk/coord.cgi?p=TR256393#llTR2560039300) | negative |  |  | 1 |  |  |
|  | positive |  |  |  |  |  |
|  | total |  |  | 1 |  |  |
| 51.3, -2.2 | negative |  |  |  | 2 |  |
|  | positive |  |  |  |  |  |
|  | total |  |  |  | 2 |  |
| Total | negative | 163 | 277 | 10 | 32 | 1 |
|  | positive | 34 |  |  |  |  |
|  | Total | 197 | 277 | 10 | 32 | 1 |

Supplementary table 2: Date of sample collection by species and E gene qPCR positivity

| Date | Sarbecovirus envelope gene qPCR results | *Rhinolophus hipposideros*  (% positive for E gene qPCR) | *Rhinolophus ferrumequinum* | *Pipistrellus* *pipistrellus* | *Myotis daubentonii* | *Myotis nattereri* | Total |
| --- | --- | --- | --- | --- | --- | --- | --- |
| not known |  |  | 9 |  |  |  | 9 |
|  |  |  | 9 |  |  |  | 9 |
| April  2021 | negative | 3 | 4 |  |  |  | 7 |
|  | Positive | 5 (65%) |  |  |  |  | 5 |
|  | total | 8 | 4 |  |  |  | 12 |
| May  2021 | negative |  | 1 |  |  |  | 1 |
|  | Positive | 4 (100%) |  |  |  |  | 4 |
|  | total | 4 | 1 |  |  |  | 5 |
| June  2021 | negative | 7 | 13 |  | 2 | 1 | 23 |
|  | Positive | 7 (50%) |  |  |  |  | 7 |
|  | total | 14 | 13 |  | 2 | 1 | 30 |
| July  2021 | negative | 4 | 6 | 4 |  |  | 14 |
|  | Positive | 1 (20%) |  |  |  |  | 1 |
|  | total | 5 | 6 | 4 |  |  | 15 |
| Aug  2021 | negative | 8 | 22 | 5 |  |  | 35 |
|  | Positive | 4 (33%) |  |  |  |  | 4 |
|  | total | 12 | 22 | 5 |  |  | 39 |
| Sept  2021 | negative | 94 | 109 |  | 28 |  | 231 |
|  | Positive | 10 (9.1%) |  |  |  |  | 10 |
|  | total | 104 | 109 |  | 28 |  | 241 |
| Oct  2021 | negative | 46 | 36 |  | 2 |  | 84 |
|  | Positive | 3 (6.5%) |  |  |  |  | 3 |
|  | total | 49 | 36 |  | 2 |  | 87 |
| Nov  2021 | negative |  | 23 |  |  |  | 23 |
|  | positive |  |  |  |  |  |  |
|  | total |  | 23 |  |  |  | 23 |
| Dec  2021 | negative |  | 1 |  |  |  | 1 |
|  | positive |  |  |  |  |  |  |
|  | total |  | 1 |  |  |  | 1 |
| Jan  2022 | negative | 1 |  | 1 |  |  | 2 |
|  | positive |  |  |  |  |  |  |
|  | total | 1 |  | 1 |  |  | 2 |
| Feb  2022 | negative |  | 53 |  |  |  | 53 |
|  | positive |  |  |  |  |  |  |
|  | total |  | 53 |  |  |  | 53 |
| Total | negative | 163 | 277 | 10 | 32 | 1 | 483 |
|  | Positive | 34 |  |  |  |  | 34 |
|  | total | 197 | 277 | 10 | 32 | 1 | 517 |

Supplementary Table 3: read statistics for sequenced samples. Raw reads and classification of reads by the Kraken2 viral database

| Sample  ID | Total Number of raw reads | Unclassified reads (%) | Classified viral reads (%) | Reads classified as *Coronaviridae* (%) |
| --- | --- | --- | --- | --- |
| RhGB02 | 25,903,657 | 25,637,600 (98.97) | 266,057 (1.03) | 1750 (0.66) |
| LH011 | 26,879,434 | 26,713,960 (99.38) | 165,474 (0.62) | 17 (0.01) |
| RhGB03 | 26,505,351 | 26,202,813 (98.86) | 302,538 (1.14) | 961 (0.32) |
| RhGB04 | 31,568,198 | 31,197,098 (98.82) | 371,100 (1.18) | 182840 (49.27) |
| RhGB05 | 32,836,967 | 32,676,343 (99.51) | 160,624 (0.49) | 102118 (63.58) |
| RhGB06 | 31,346,436 | 31,109,734 (99.24) | 236,702 (0.76) | 134323 (56.75) |

Supplementary Table 4: genome length, quality, average amino acid identity with reference genome assembly, completeness and contamination following CheckV genome assessment.

| Sample ID | Length (kb) | Quality (%) | Average amino acid identity | Completeness (%) | Contamination (%) |
| --- | --- | --- | --- | --- | --- |
| RhGB02 | 29.5 | 98 | 84.98 | 97.7 |  |
| RhGB03 | 28.2 | 94 | 84.05 | 94.03 |  |
| RhGB04 | 12.6 | 42 | 87.95 | 42.14* |  |
| RhGB05 | 30.6 | 96 | 84.92 | 96.94 | 3.17 |
| RhGB06 | 30. 4 | 100 | 84.97 | 100 |  |

Supplementary Table 5: Details on the number of reads, coverage, and mean depth, and reads mapping quality assessment on lesser horseshoe bat sarbecovirus reported from this study. Kb (kilobase), % (percentage), * (incomplete genome).

| Sample ID | Start position | End position | Number of reads | Coverage bases | Coverage (%) | Mean depth | Mean base quality | mean map quality |
| --- | --- | --- | --- | --- | --- | --- | --- | --- |
| RhGB02 | 1 | 29566 | 7148 | 29566 | 100 | 35.7409 | 35.7 | 59.4 |
| RhGB03 | 1 | 28214 | 4264 | 28214 | 100 | 22.434 | 35.5 | 59.7 |
| RhGB04 | 1 | 12647 | 302390 | 12647 | 100 | 3446.35 | 36.1 | 59.1 |
| RhGB05 | 1 | 30600 | 420416 | 30600 | 100 | 2026.71 | 35.6 | 59.4 |
| RhGB06 | 1 | 30463 | 593793 | 30463 | 100 | 2867.89 | 35.8 | 59.4 |

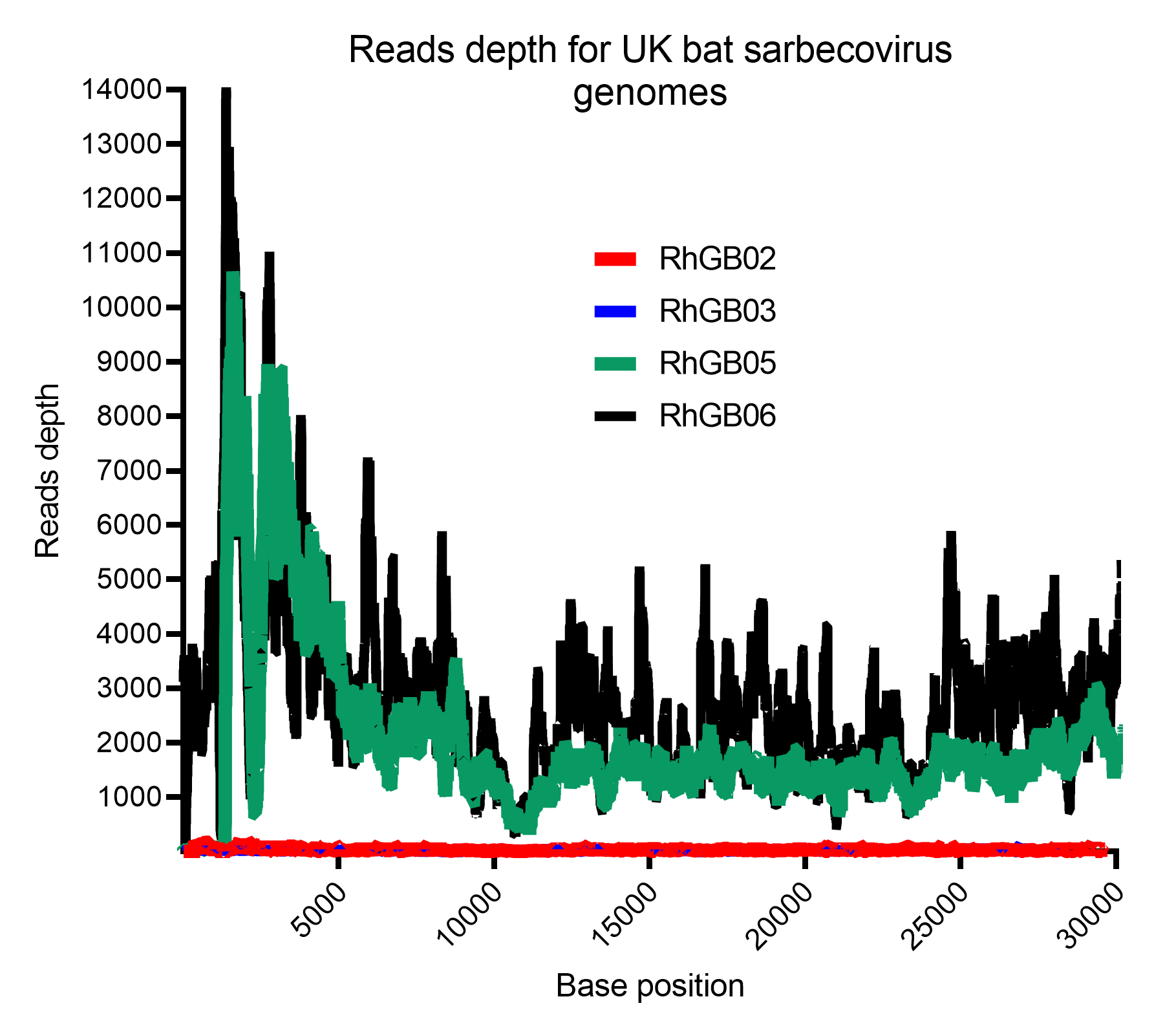

Supplementary figure 1 : Read depth for each near full length coronavirus contig recovered from lesser horseshoe bats.

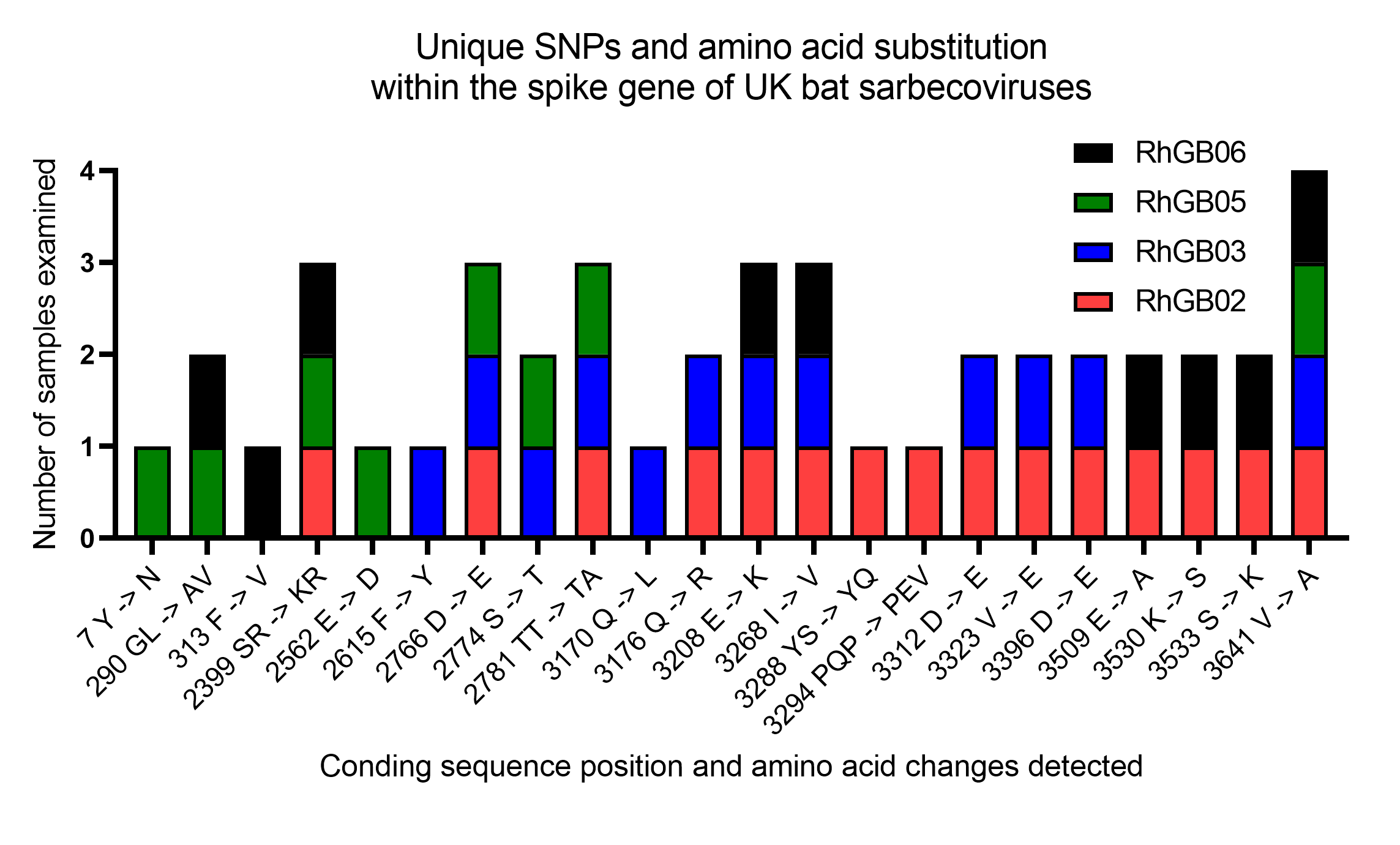

Supplementary Figure 2: Unique SNPs and amino acid substitutions reported within the spike gene of UK bat sarbecovirus assembled genomes following reads mapping to the European lesser horseshoe bat sarbecovirus reference genome Bat CoV BM48-31 NC01170 and variant calling. The spike glycoprotein of all assembled UK lesser horseshoe bat sarbecovirus consisted of 3771 bp, while that of the Bulgarian sarbecovirus reference sequence had 3780 bp. The numbers displayed behind amino acid changes along the X-axis include nucleotide position beginning at position 1 to 3771 bp. Below this the chart is a linear map of the UK bat sarbecovirus spike glycoprotein sequence showing the presence of multiple SNPs within the SD-1 and SD-2 subdomains, S1/S2 cleavage region and the S2 fusion subunit of the spike glycoprotein.

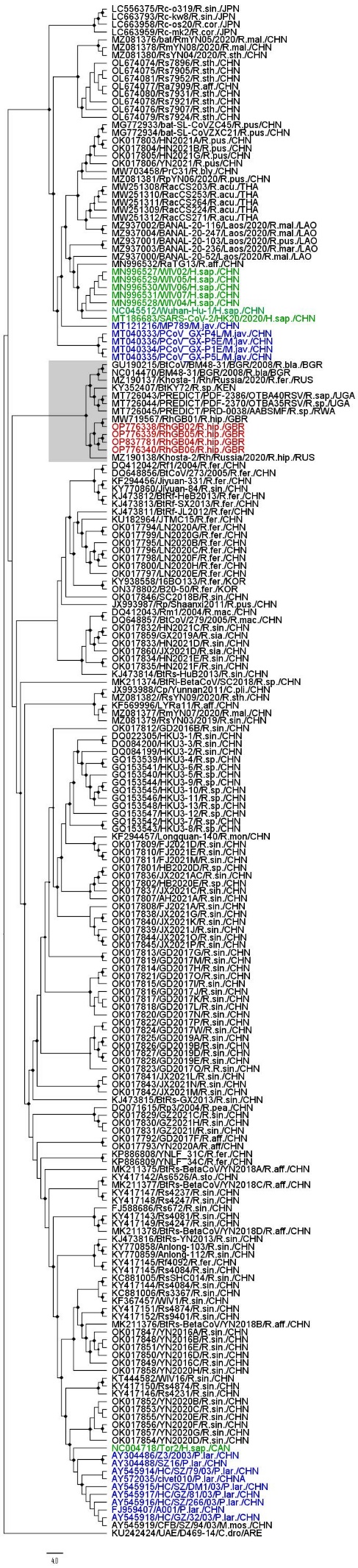

Supplementary Figure 3: Maximum likelihood phylogenetic tree full genome nucleic acid tree constructed with 1000 bootstrap approximation, rooted on the MERs coronavirus reference sequence. One hundred and ninety eight non-human sarbecovirus genomes and reference sequences for major variants of SARS-CoV and SARS-CoV-2 were included (non-human SARS-CoV-2 isolates were not included). Red= isolates from this study, Green=human isolates , blue=isolates from other mammals. Sequences are named with Genbank ID, name from original study species of origin (eg R.hip= Rhinolophus hipposiderus) and country of origin (eg GBR= Great Britain)

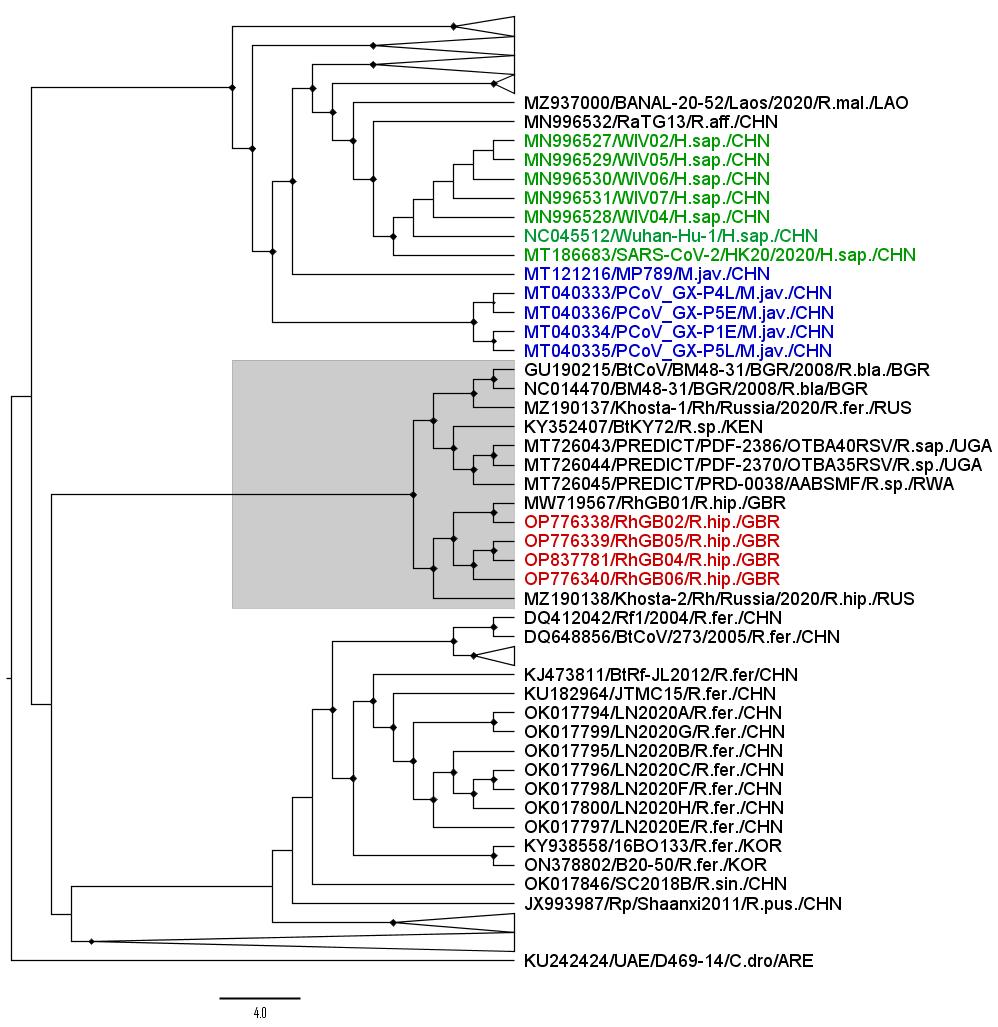

Supplementary Figure 4: Maximum likelihood phylogenetic tree full genome nucleic acid tree constructed with 1000 bootstrap approximation, rooted on the MERs coronavirus reference sequence. One hundred and ninety eight non-human sarbecovirus genomes and reference sequences for major variants of SARS-CoV and SARS-CoV-2 were included (non-human SARS-CoV-2 isolates were not included) . Clades of Asian Bat coronavirus sequences (apart from greater horseshoe bat sequences) have been collapsed for clarity (represented as broad triangles). Red= isolates from this study, Green=human isolates , blue=isolates from other mammals. Sequences are named with Genbank ID, name from original study species of origin (eg R.hip= Rhinolophus hipposiderus) and country of origin (eg GBR= Great Britain)

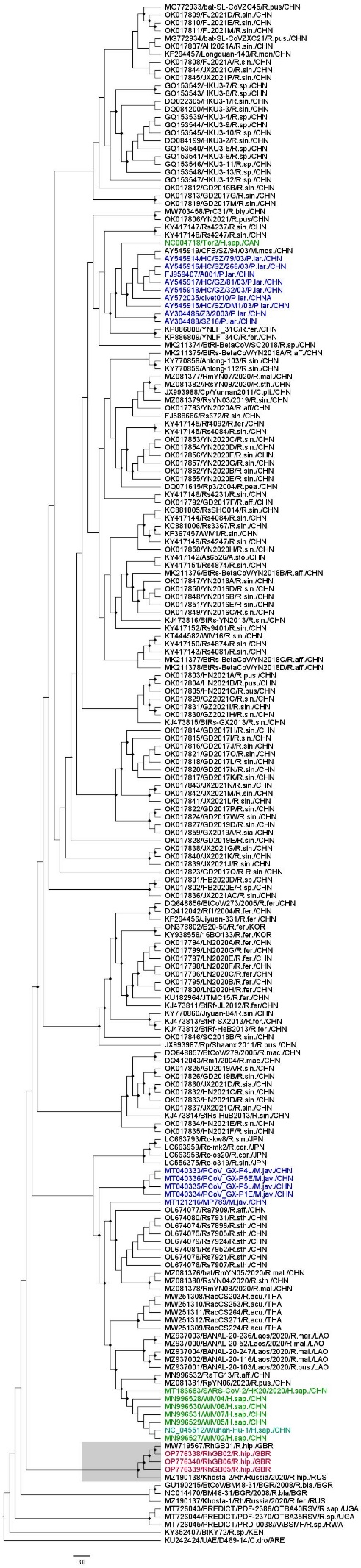

Supplementary Figure 5: Maximum likelihood phylogenetic tree RDRP gene nucleic acid tree constructed with 1000 bootstrap approximation, rooted on the MERs coronavirus reference sequence. One hundred and ninety eight non-human sarbecovirus genomes and reference sequences for major variants of SARS-CoV and SARS-CoV-2 were included (non-human SARS-CoV-2 isolates were not included) . Red= isolates from this study, Green=human isolates , blue=isolates from other mammals. Sequences are named with Genbank ID, name from original study species of origin (eg R.hip= Rhinolophus hipposiderus) and country of origin (eg GBR= Great Britain)

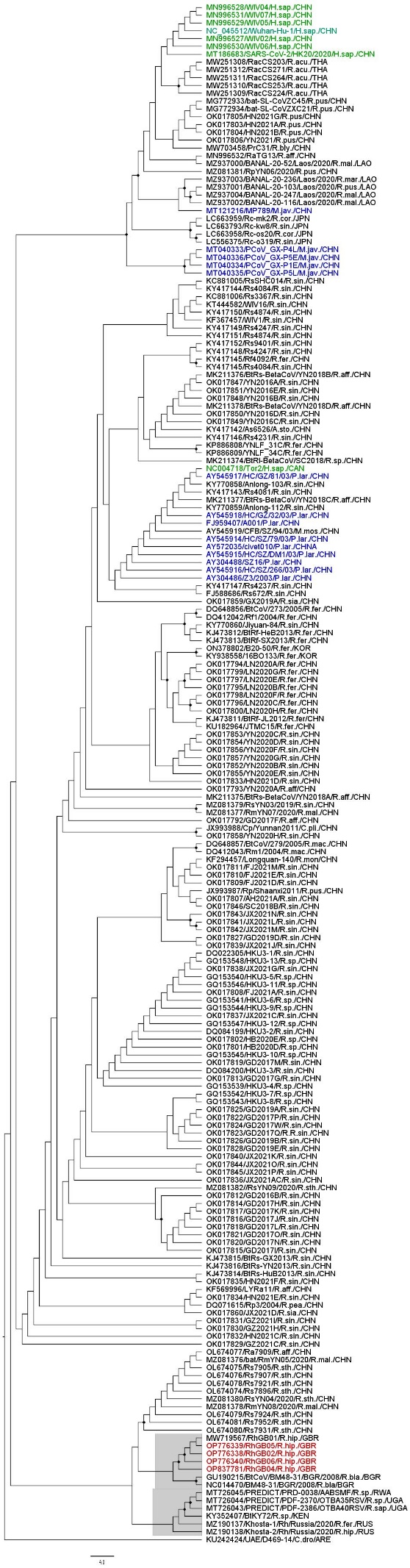

Supplementary Figure 6: Maximum likelihood phylogenetic tree E gene nucleic acid tree constructed with 1000 bootstrap approximation, rooted on the MERs coronavirus reference sequence. One hundred and ninety eight non-human sarbecovirus genomes and reference sequences for major variants of SARS-CoV and SARS-CoV-2 were included (non-human SARS-CoV-2 isolates were not included) . Red= isolates from this study, Green=human isolates , blue=isolates from other mammals. Sequences are named with Genbank ID, name from original study species of origin (eg R.hip= Rhinolophus hipposiderus) and country of origin (eg GBR= Great Britain)

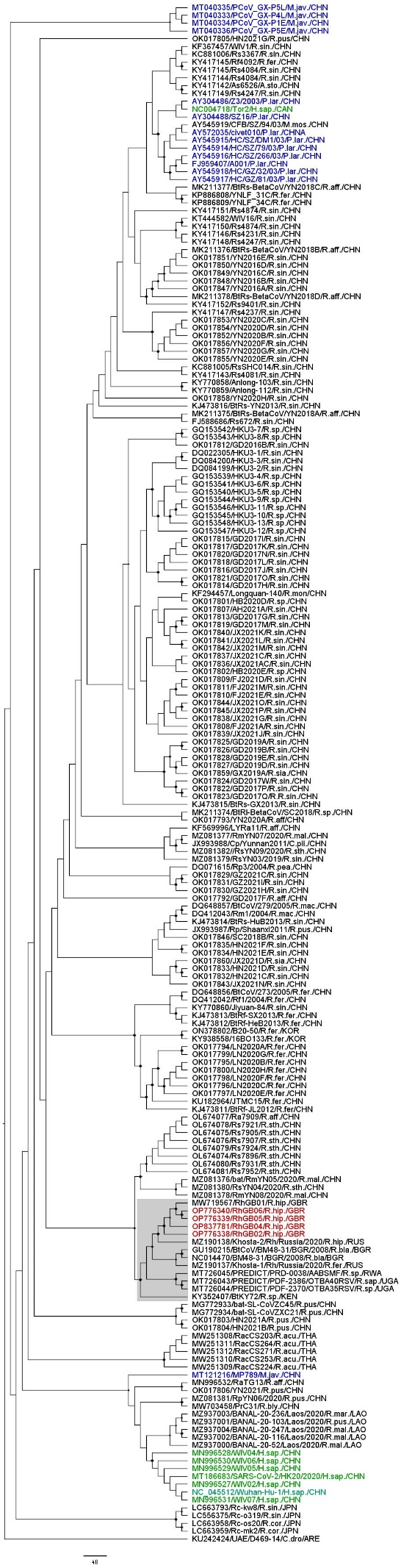

Supplementary Figure 7: Maximum likelihood phylogenetic tree N gene nucleic acid tree constructed with 1000 bootstrap approximation, rooted on the MERs coronavirus reference sequence. One hundred and ninety eight non-human sarbecovirus genomes and reference sequences for major variants of SARS-CoV and SARS-CoV-2 were included (non-human SARS-CoV-2 isolates were not included) . Red= isolates from this study, Green=human isolates , blue=isolates from other mammals. Sequences are named with Genbank ID, name from original study species of origin (eg R.hip= Rhinolophus hipposiderus) and country of origin (eg GBR= Great Britain)

| UK bat CoVs (RhGB05) gene/proteins | UK bat CoVs reported from this study | | | | | Bat CoV RhGB01 MW719567 | Bat CoV BM48-31 NC014470 | SARS-CoV NC004718 | SARS-CoV-2 NC045512 |
| --- | --- | --- | --- | --- | --- | --- | --- | --- | --- |
|  | RhGB05 aa  length | RhGB02 | RhGB03 | RhGB04 | RhGB06 |  |  |  |  |
| ORF1ab (nsp1 ─ 16) | Amino acid Percentage identity | | | | | | | | |
| nsp1 | 180 | 98.9 | 100 | N/A | 100 | 98.9 | 85.6 | 77.8 | 80.6 |
| nsp2 | 638 | 96.2 | 99.8 | N/A | 98.1 | 96.1 | 74.3 | 68.2 | 63.8 |
| nsp3 | 1908 | 98.2 | 99.2 |  | 98.4 | 98.1 | 75.6 | 73.5 | 69.5 |
| nsp4 | 500 | 98.6 | 100 |  | 99.6 | 98.8 | 81.6 | 82.2 | 79.2 |
| nsp5 | 306 | 99 | 100 |  | 99.7 | 99 | 90.2 | 93.5 | 91.2 |
| nsp6 | 290 | 98.3 | 98.6 |  | 98.6 | 98.3 | 86.9 | 87.2 | 84.8 |
| nsp7 | 83 | 100 | 100 |  | 100 | 100 | 92.8 | 92.8 | 94 |
| nsp8 | 198 | 99.5 | 100 |  | 100 | 100 | 93.9 | 94.4 | 92.4 |
| nsp9 | 113 | 100 | 100 |  | 100 | 100 | 92.9 | 93.8 | 92 |
| nsp10 | 139 | 100 | 100 |  | 99.3 | 100 | 92.8 | 94.2 | 93.5 |
| nsp11 | 13 | 100 | 92.3 |  | 100 | 15.4 | 84.6 | 30.8 | 23.1 |
| nsp12 | 923 | 99.3 | 99.7 |  | 99.7 | 14 | 11.7 | 96.1 | 11.4 |
| nsp13 | 601 | 99.3 | 99.3 | 14.3 | 99.5 | 99.2 | 94.7 | 94.3 | 94.5 |
| nsp14 | 527 | 99.2 | 99.2 | 100 | 99.2 | 99.2 | 93.7 | 94.3 | 92.6 |
| nsp15 | 346 | 97.7 | 22.6 | 99.1 | 99.1 | 97.7 | 90.8 | 92.5 | 90.5 |
| nsp16 | 298 | 99.7 | 98.7 | 99.7 | 99.3 | 99.7 | 90.9 | 90.9 | 88.9 |
| S | 1256 | 97.9 | 97.7 | 98.2 | 97.9 | 98.4 | 81.3 | 72.7 | 71.2 |
| ORF3a | 271 | 98.9 | 97.8 | 99.6 | 100 | 98.5 | 79.4 | 69.5 | 65.9 |
| ORF3b | 150 | 96 | 96 | 97.3 | 99.3 | 97.3 | 56.3 | 43.9 | 40.4 |
| E | 77 | 100 | 100 | 100 | 100 | 100 | 90.9 | 94.8 | 90.9 |
| M | 223 | 100 | 100 | 99.6 | 100 | 99.6 | 90.8 | 91.9 | 88.8 |
| ORF6a | 116 | 98.3 | 97.4 | 96.6 | 99.1 | 96.6 | 60.3 | 47.9 | 38.5 |
| ORF6b | 63 | 98.4 | 95.2 | 93.7 | 98.4 | 93.7 | 60.3 | 50 | 43.5 |
| ORF7a | 119 | 96.6 | 95.8 | 98.3 | 95.8 | 95.8 | 55.8 | 54.5 | 51.2 |
| ORF7b | 43 | N/A | 100 | 100 | N/A | 100 | 78 | 64.3 | 64.3 |
| ORF9b | 97 | 99 | N/A | 96.9 | 96.9 | 99 | 75.3 | 48.5 | 48 |
| N | 418 | 99.5 | 10.3 | 99 | 99.5 | 99.8 | 91.4 | 85.8 | 85.5 |
| ORF10 | 39 | 100 | N/A | N/A | 100 | 100 | 100 | 97.4 | 79.5 |

**Supplementary table 6**: A comparison of protein composition and percentage identity for structural, non-structural, and accessory proteins between UK bat CoVs (RhGB05, RhGB02, RhGB03, RhGB05, and RhGB06), related bat SARS-like virus, SARS-CoV, and SARS-CoV-2 reference CoVs. For clarity only one of the UK bats CoVs assembled genomes was used to compared amino acid percentage identity against related and reference genomes.

Accessible

A

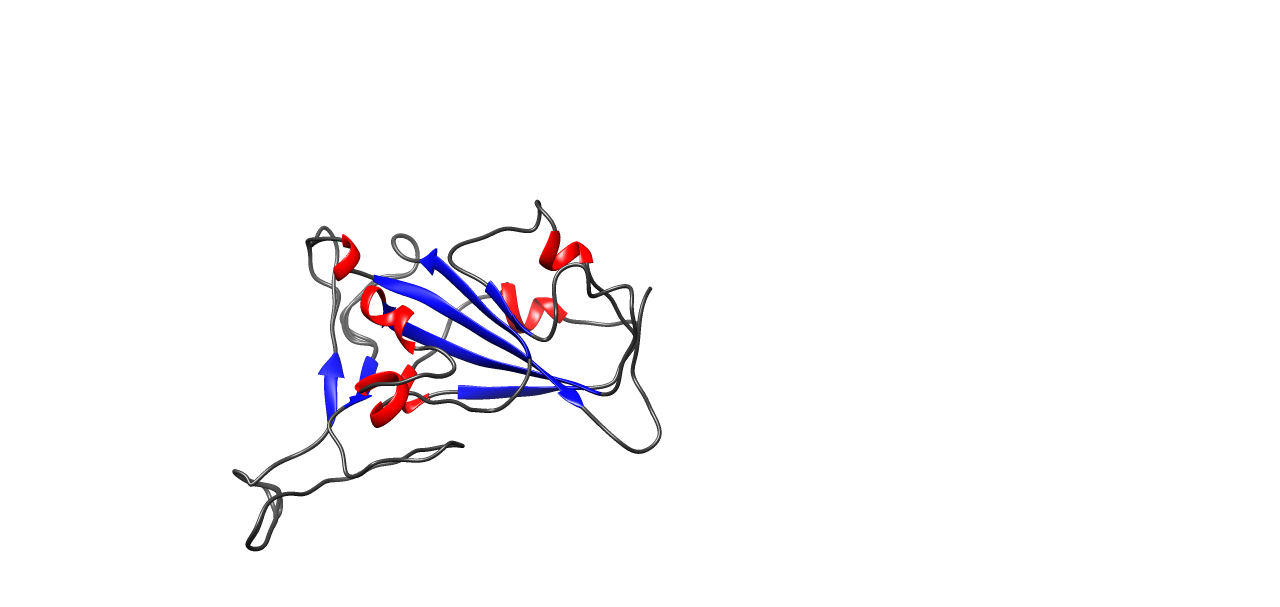

B

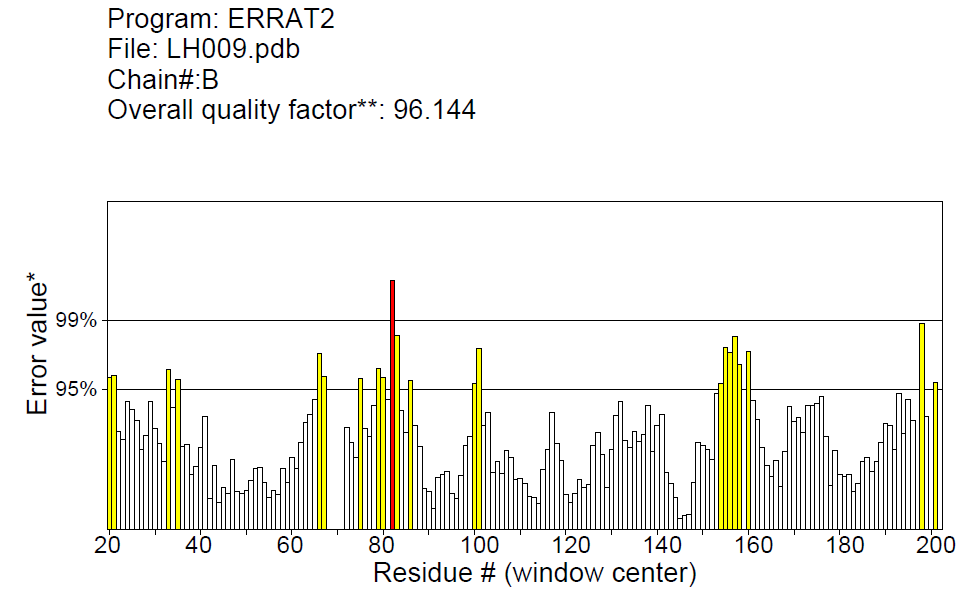

C

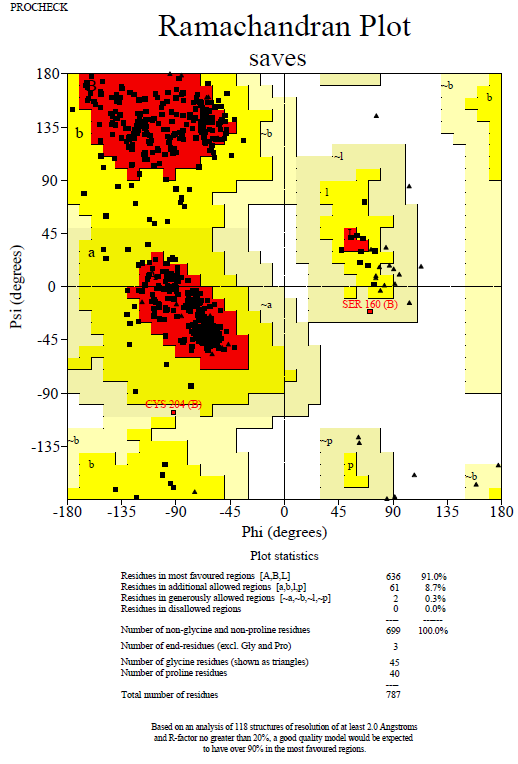

Supplementary figure 8: (A) Represents 3D models helices (orange), stands (blue) and coils (dark gray), RBD 3D models were visualized and superimposed from five UK bat CoVs (RhGB02 (renamed from LH009), RhGB03, RhGB04 and RhGB06) using Chimera. (B) Represents bar charts generated following evaluation of models constructed using ERRAT2 with the y-axis representing 95% (yellow bars) and 99% (red bars) confidence rejection level for the segment of RBD proteins analyzed. (C) shows a Ramachandran plot generated from 3D validation showing RBD amino acid residues generously allowed (light yellow), allowed (yellow) and favoured (red) during 3D complex modelling
